# Rapid Diagnosis of Lower Respiratory Infection using Nanopore-based Clinical Metagenomics

**DOI:** 10.1101/387548

**Authors:** Themoula Charalampous, Hollian Richardson, Gemma L. Kay, Rossella Baldan, Christopher Jeanes, Duncan Rae, Sara Grundy, Daniel J. Turner, John Wain, Richard M. Leggett, David M. Livermore, Justin O’Grady

## Abstract

Lower respiratory infections (LRIs) accounted for three million deaths worldwide in 2016, the leading infectious cause of mortality. The “gold standard” for investigation of bacterial LRIs is culture, which has poor sensitivity and is too slow to guide early antibiotic therapy. Metagenomic sequencing potentially could replace culture, providing rapid, sensitive and comprehensive results. We developed a metagenomics pipeline for the investigation of bacterial LRIs using saponin-based host DNA depletion combined with rapid nanopore sequencing. The first iteration of the pipeline was tested on respiratory samples from 40 patients. It was then refined to reduce turnaround and increase sensitivity, before testing a further 41 samples. The refined method was 96.6% concordant with culture for detection of pathogens and could accurately detect resistance genes with a turnaround time of six hours. This study demonstrates that nanopore metagenomics can rapidly and accurately characterise bacterial LRIs when combined with efficient human DNA depletion.

---

Lower respiratory infections (LRIs) are the third most common cause of death globally and are the leading infectious cause, accounting for three million deaths worldwide in 2016 (http://www.who.int/news-room/fact-sheets/detail/the-top-10-causes-of-death). They can be subdivided into chest infection, community-acquired pneumonia (CAP), hospital-acquired pneumonia (HAP), bronchitis and bronchiolitis (https://www.nice.org.uk/guidance/cg191). Morbidity and mortality rates vary dependent on infection site, pathogen and host factors. In the UK, CAP accounts for approx. 29,000 deaths per annum and it is estimated that HAP accounts for approx. 36,000 deaths in the US per annum ^1^. The most common bacterial agents of LRIs are *Streptococcus pneumoniae* and *Haemophilus influenzae* for CAP and *Staphylococcus aureus*, Enterobacteriaceae and *Pseudomonas aeruginosa* for HAP ^2–6^. However, a wide range of pathogens, including viruses, can cause these infections, meaning that microbial diagnosis and treatment are challenging.

Every year in the UK NHS treat 16 million people with antibiotics for respiratory tract infections (https://www.nice.org.uk/guidance/cg191/documents/pneumonia-final-scope2). Initial treatment for severe LRIs, particularly HAP, often involves empirical broad-spectrum antibiotics. Guidelines recommend that such therapy should be refined or stopped after two to three days, once microbiology results become available ^7–9^, but this is often not done, if the patient is responding well or the laboratory has failed to identify a pathogen. Such extensive ‘blind’ use of broad-spectrum antibiotics is wasteful and constitutes poor stewardship, given that many patients are infected with a virus or a susceptible pathogen. Antimicrobial over-treatment leads to disruption of the gut flora, promoting the risk of over-growth by resistant bacteria and of *Clostridium difficile*^10, 11^.

There is an urgent need for rapid microbiological diagnostics to ensure swift tailored treatment and to reduce prolonged broad-spectrum antibiotic therapy. “Gold standard” culture and susceptibility testing is not fit for purpose in this regard, owing to slow turnaround times (48-72 hours) and low clinical sensitivity ^1, 2, 7, 12^. Molecular methods have the potential to overcome this limitations, as highlighted by the Chief Medical Officer and O'Neill reports (2016), identifing pathogens and their antibiotic resistance profiles within a few hours, thereby allowing early therapeutic refinement, and supporting effective antibiotic stewardship ^13,14^. However, molecular methods present their own challenges. Nucleic acid amplification tests (including PCR) are rapid and highly specific/sensitive, but uncommon agents and antibiotic resistances cannot readily be sough, given the limits on multiplexing ^8, 15, 16^. ^17^. There is also a constant need to update these PCR-based methods to include emerging resistance genes and mutations^9, 18, 19^.

Metagenomic sequencing based approaches, which make no presumptions about the organisms and resistance genes that may be present, have the potential to overcome the shortcomings of both culture and PCR, by combining speed with comprehensiveness ^20^. Nanopore sequencing (Oxford Nanopore Technologies, ONT) has particular potential for rapid diagnostic applications, with the advantage of real-time data acquisition and analysis ^21, 22^ as compared with other NGS platforms (e.g. Illumina) where sequencing can take up to 48hrs before analysis can begin ^23^. Nanopore sequencing has previously been used to identify viral and bacterial pathogens from clinical samples using targeted approaches and in proof-of-concept studies using samples with high pathogen loads (e.g. urinary tract infection) ^22, 24, 25^. Respiratory specimens present a greater challenge, owing to the variable pathogen load, the presence of commensal respiratory tract flora, and the high ratio of host:pathogen nucleic acids present (up to 10^5^:1 in sputum).

It is possible to apply rapid metagenomics to respiratory samples, as demonstrated by Pendleton *et al*. (2017), who used nanopore sequencing, without host cell depletion, to samples from two bacterial pneumonia patients. The vast majority of reads, however, were of human origin, with only one and two nanopore reads aligned to the infecting pathogens, *P. aeruginosa* and *S. aureus*, respectively ^26^. It seems likely that their method could be improved by depletion of host DNA but, although commercial kits and published methods are available for this purpose ^27, 28^, they do not perform well in complex respiratory samples and improved methods are required.

Accordingly, we developed and optimised a state-of-the-art nanopore-sequencing based clinical metagenomics pipeline for the investigation of bacterial LRIs capable of removing up to 99.99% host nucleic acid from clinical respiratory samples, enabling pathogen and antibiotic resistance gene identification within six hours.

## RESULTS

### Pilot method testing

The first ‘Pilot’ iteration of the metagenomics pipeline, the pilot method, was tested on respiratory samples from 40 patients with suspected bacterial LRI. Overall, this method was 91.2% sensitive (95% CI; 75.2-97.7%) and 83.3% specific (95% CI; 36.5-99.1%), *not* counting additional organisms in culture-positive samples as false positives (Table 1), and took approx. 8 hours to perform (Figure 1). Up to 99.9% or ~10^3^ fold (average: 10^3^ fold; range: 32-1024 fold) of host DNA was removed using the saponin depletion. Pathogens were identified in real-time using ONT’s What’s In My Pot? (WIMP) pipeline. Additional pathogenic bacteria were detected in 6/40 samples: *S. pneumoniae* was detected in P20; *Moraxella catarrhalis* in P8; *Escherichia coli* in P14; *H. influenzae* in P22 and P30; *Klebsiella pneumonia*e and *M. catarrhalis* in P29 (Supplementary Table 1).

**Table 1:** Pilot metagenomic pipeline output compared to routine microbiology culture results.

| Pathogen | Culture positive | Metagenomics |  |  |
| --- | --- | --- | --- | --- |
|  |  | Detected | Missed detection | Additional detection |
| *Haemophilus influenzae* | 10 | 9 | 1 | 2 |
| *Staphylococcus aureus* | 8 | 7 | 1 | 0 |
| Coliform* | 6 | 6** | 0 | 0 |
| *Streptococcus pneumoniae* | 6 | 5 | 1 | 1 (1***) |
| *Klebsiella pneumoniae* | 3 | 3 | 0 | 1 |
| *Pseudomonas aeruginosa* | 3 | 3 | 0 | 0 |
| *Enterobacter aerogenes* | 1 | 1 | 0 | 0 |
| *Enterobacter cloacae* complex | 1 | 1 | 0 | 0 |
| *Escherichia coli* | 1 | 1 | 0 | 1 |
| *Moraxella catarrhalis* | 1 | 1 | 0 | 2 |
| None (NRF/NSG) | 0 | 0 | 0 | 1 |

\*Coliform not further identified by culture \*\* 2 x *Escherichia coli*, *Klebsiella oxytoca*, *Klebsiella pneumoniae*, *Proteus mirabilis*, *Serratia marcescens* \*\*\*Additional detection in NRF/NSG samples

**Figure 1:**
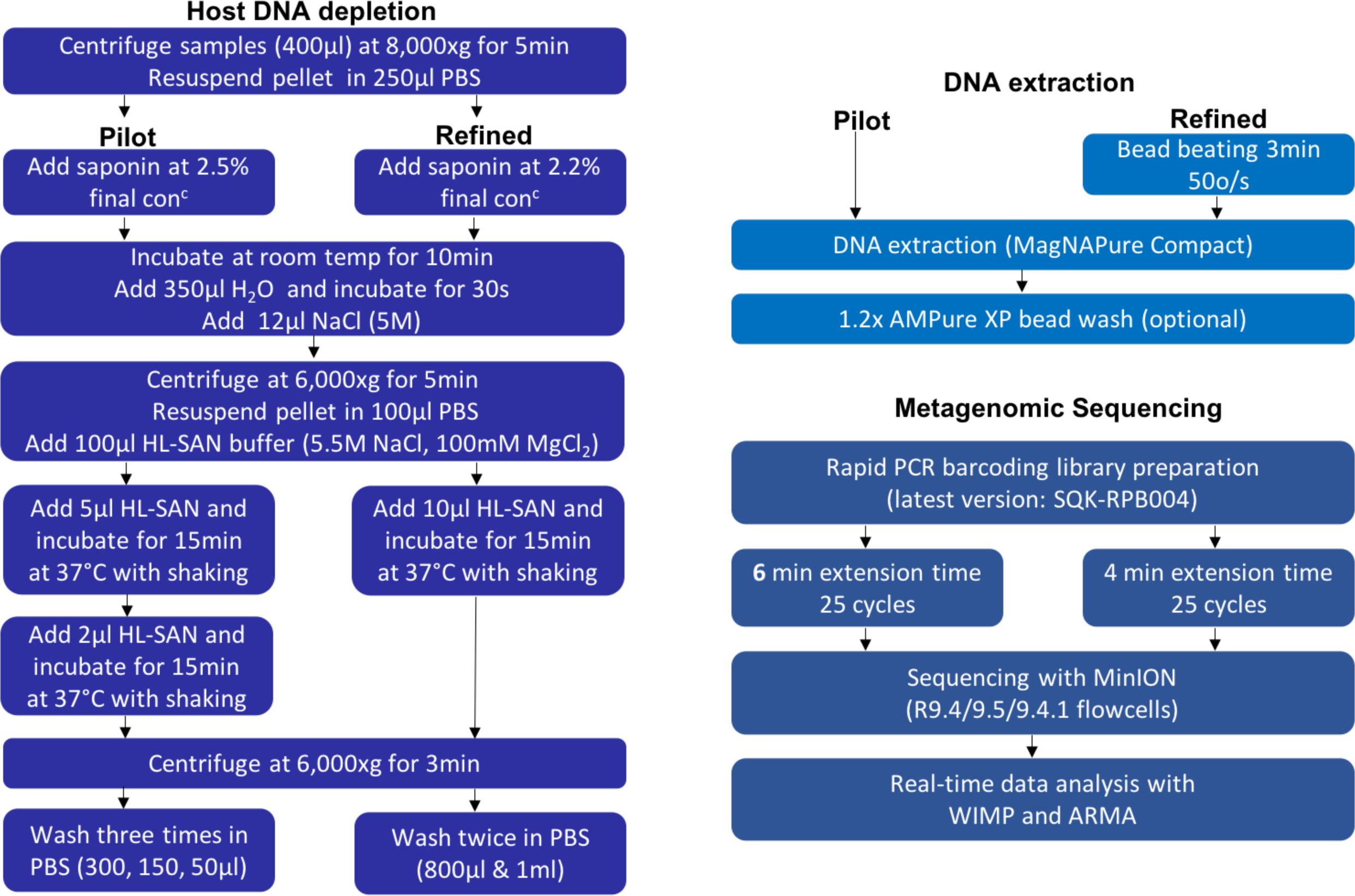
Schematic representation of the metagenomic pipeline with a turnaround time of approx. six hours (refined) and approx. eight hours (pilot) from sample collection to sample result.

Organisms reported by routine microbiology were not detected in 3/40 samples by the Pilot method. Two of these cases were mixed infections where one organism was missed by the Pilot method – specifically, *S. pneumoniae* and *H. influenzae* were not detected in P3 and P37 respectively - and *S. aureus* was not detected in P34.

### Optimisation experiments

Efforts were made to reduce major errors (8.8% false negative rate) by improving bacterial cell lysis, thereby improving assay performance. We also sought to reduce the sample-to-result turnaround time.

#### Improving bacterial cell lysis

A sample pre-treatment step was introduced, involving either bead-beating or an enzyme cocktail, to increase the lysis of difficult-to-lyse organisms. Two culture-positive sputa were used for optimisation experiments, one containing *S. aureus* (Gram-positive) and one containing *P. aeruginosa* (Gram-negative). Neither pre-treatment affected the yield of *P. aeruginosa* DNA, but the enzyme cocktail increased the amount of *S. aureus* DNA detected by approx. 4-fold and bead-beating achieved a 20-fold increase compared with the pilot method, as determined by qPCR (Supplementary Table 2a). Accordingly, bead-beating was introduced as a standard pre-treatment in the Optimised method.

**Table 2:** Human and bacterial DNA qPCR results for sputum samples infected by Gram-negative and Gram-positive bacteria with and without host nucleic acid depletion

| Sample | Sample type | Organism cultured by microbiology<br>(Organism identified from metagenomic pipeline) | Sample treatment | Human qPCR assay (Cq) | Human DNA depletion (ΔCq) | 16S rRNA gene V3-V4 fragment qPCR assay (Cq) | Bacterial gain/loss to standard depletion (ΔCq) |
| --- | --- | --- | --- | --- | --- | --- | --- |
| S1 | ETA | *E. coli*<br>(*E. coli*) | Undepleted | 22.62 | 12.38 | 15.60 | 0.13 |
| | | | Depleted | 35.00 | ($\sim 10^4$) | 15.73 | |
| S2 | Sputum | *K. pneumoniae*<br>(*K. pneumoniae*) | Undepleted | 23.73 | 9.99 | 15.63 | 0.02 |
| | | | Depleted | 33.71 | ($\sim 10^3$) | 15.65 | |
| S3 | Sputum | *P. aeruginosa*<br>(*P. aeruginosa*) | Undepleted | 23.05 | 9.29 | 15.46 | 1.48 |
| | | | Depleted | 32.34 | ($\sim 10^3$) | 13.98 | |
| S4 | Sputum | *S. mariscens*<br>(*S. mariscens*) | Undepleted | 26.34 | 9.93 | 16.96 | 0.52 |
| | | | Depleted | 36.27 | ($\sim 10^3$) | 17.48 | |
| S5 | Sputum | *K. oxytoca*<br>(*K. oxytoca* & *K. pneumoniae*) | Undepleted | 22.96 | 8.58 | 12.67 | 0.64 |
| | | | Depleted | 31.54 | ($\sim 10^3$) | 12.03 | |
| S6 | Sputum | *S. aureus*<br>(*S. aureus*) | Undepleted | 22.31 | 9.41 | 19.11 | 1.57 |
| | | | Depleted | 31.72 | ($\sim 10^3$) | 17.54 | |
| S7 | Sputum | *H. influenzae*<br>(*H. influenzae* & *P. aeruginosa*) | Undepleted | 25.47 | 9.53 | 21.44 | 0.43 |
| | | | Depleted | 35.00 | ($\sim 10^3$) | 21.87 | |
| S8 | Sputum | *M. catarrhalis*<br>(*M. catarrhalis* & *S. pneumoniae*) | Undepleted | 22.72 | 9.17 | 16.9 | 0.66 |
| | | | Depleted | 31.89 | ($\sim 10^3$) | 17.56 | |
| S9 | Sputum | *P. aeruginosa* & *E. coli*<br>(*E. coli*) | Undepleted | 23.89 | 11.11 | 19.58 | 3.26 |
| | | | Depleted | 35 | ($\sim 10^4$) | 22.84 | |
| S10 | Sputum | *H. influenzae*<br>(*H. influenzae* & *S. pneumoniae*) | Undepleted | 23.46 | 8.6 | 14.12 | 2.39 |
| | | | Depleted | 32.06 | ($\sim 10^3$) | 16.51 | |
| S11 | Sputum | NRF<br>(*S. pneumoniae*) | Undepleted | 25.77 | 9.23 | 17.96 | 1.92 |
| | | | Depleted | 35.00 | ($\sim 10^3$) | 19.88 | |
| S12 | Sputum | NRF<br>(*H. influenzae* & *M. catarrhalis*) | Undepleted | 22.5 | 8.92 | 17.61 | 0.05 |
| | | | Depleted | 31.42 | ($\sim 10^3$) | 17.56 | |
| S13 | Sputum | *S. mariscens*<br>(*S. mariscens*) | Undepleted | 22.48 | 7.11 | 12.77 | 0.79 |
| | | | Depleted | 29.59 | ($\sim 10^2$) | 11.98 | |
| **S14** | Sputum | *S. aureus*<br>(*S. aureus* & *M. catarrhalis*) | Undepleted | 23.17 | 7.68 | 13.83 | 0.96 |
|  |  |  | Depleted | 30.85 | (~10<sup>2</sup>) | 14.79 |  |
| **S15** | Sputum | *S. aureus*<br>(*S. aureus* & *S. pneumoniae*) | Undepleted | 22.66 | 8.47 | 18.73 | 0.08 |
|  |  |  | Depleted | 31.13 | (~10<sup>3</sup>) | 18.65 |  |
| **S16** | Sputum | MRSA<br>(MRSA) | Undepleted | 25.51 | 6.43 | 15.32 | 0.24 |
|  |  |  | Depleted | 31.94 | (~10<sup>2</sup>) | 15.56 |  |
| **S17** | Sputum | NRF<br>(None) | Undepleted | 23.51 | 9.64 | 19.55 | 1.17 |
|  |  |  | Depleted | 33.15 | (~10<sup>3</sup>) | 20.72 |  |
| **S18** | Sputum | *H. influenzae*<br>(*H. influenzae*) | Undepleted | 27.14 | 7.86 | 12.89 | 2.21 |
|  |  |  | Depleted | 35.00 | (~10<sup>2</sup>) | 15.10 |  |
| **S19** | Sputum | NRF<br>(None) | Undepleted | 22.63 | 11.18 | 19.69 | 0.69 |
|  |  |  | Depleted | 33.81 | (~10<sup>3</sup>) | 19.00 |  |
| **S20** | Sputum | *H. influenzae*<br>(*H. influenzae*) | Undepleted | 22.44 | 10.03 | 14.99 | 1.19 |
|  |  |  | Depleted | 32.47 | (~10<sup>3</sup>) | 16.18 |  |
| **S21** | Sputum | NRF<br>(*H. influenzae* & *S. pneumoniae*) | Undepleted | 24.58 | 10.42 | 16.60 | 0.82 |
|  |  |  | Depleted | 35.00 | (~10<sup>3</sup>) | 17.42 |  |
| **S22** | Sputum | NRF<br>(None) | Undepleted | 22.71 | 9.22 | 14.62 | 0.39 |
|  |  |  | Depleted | 31.93 | (~10<sup>3</sup>) | 15.01 |  |
| **S23** | Sputum | *H. influenzae*<br>(*H. influenzae*) | Undepleted | 24.82 | 10.18 | 16.80 | 1.84 |
|  |  |  | Depleted | 35.00 | (~10<sup>3</sup>) | 18.64 |  |
| **S24** | Sputum | *H. influenzae*<br>(*H. influenzae*) | Undepleted | 22.24 | 10.17 | 15.70 | 1.63 |
|  |  |  | Depleted | 32.41 | (~10<sup>3</sup>) | 17.33 |  |
| **S25** | Sputum | *H. influenzae*<br>(*H. influenzae*) | Undepleted | 25.52 | 6.26 | 16.59 | 2.67 |
|  |  |  | Depleted | 31.79 | (~10<sup>2</sup>) | 19.26 |  |
| **S26** | Sputum | *M. catarrhalis*<br>(*M. catarrhalis*) | Undepleted | 23.47 | 11.53 | 19.26 | 0.74 |
|  |  |  | Depleted | 35.00 | (~10<sup>4</sup>) | 20.00 |  |
| **S27** | Sputum | *H. influenzae* & *S. aureus*<br>(*H. influenzae* & *S. aureus*) | Undepleted | 32.74 | 2.26 | 23.19 | 7.92 |
|  |  |  | Depleted | 35.00 | (~5) | 15.27 |  |
| **S28** | Sputum | NRF<br>(*S. pneumoniae*) | Undepleted | 24.46 | 10.54 | 22.28 | 2.80 |
|  |  |  | Depleted | 35.00 | (~10<sup>3</sup>) | 25.08 |  |
| **S29** | Sputum | *P. aeruginosa*<br>(*P. aeruginosa* & *S. aureus*) | Undepleted | 24.05 | 5.11 | 19.81 | 2.04 |
|  |  |  | Depleted | 29.13 | (~10<sup>2</sup>) | 17.77 |  |
| **S30** | BAL | *P. aeruginosa*<br>(*P. aeruginosa*) | Undepleted | 29.93 | 5.07 | 22.68 | 0.00 |
|  |  |  | Depleted | >35.00 | (~33) | 22.68 |  |
| **S31** | Sputum | NRF<br>(*H. influenzae*) | Undepleted | 21.57 | 8.26<br>(~10<sup>3</sup>) | 19.79 | 1.65 |
|  |  |  | Depleted | 29.83 |  | 21.44 |  |
| **S32** | Sputum | NSG<br>(*E. coli*) | Undepleted | 25.56 | 8.68<br>(~10<sup>3</sup>) | 15.98 | 0.47 |
|  |  |  | Depleted | 34.24 |  | 16.45 |  |
| **S33** | Sputum | NRF<br>(None) | Undepleted | 21.73 | 10.04<br>(~10<sup>3</sup>) | 20.69 | 0.81 |
|  |  |  | Depleted | 31.77 |  | 21.50 |  |
| **S34** | Sputum | NSG<br>(None) | Undepleted | 25.17 | 5.40<br>(~10<sup>2</sup>) | 22.92 | 0.01 |
|  |  |  | Depleted | 30.57 |  | 22.93 |  |
| **S35** | Sputum | *E. coli*<br>(*E. coli*) | Undepleted | 21.11 | 5.18<br>(~10<sup>2</sup>) | 16.49 | 0.58 |
|  |  |  | Depleted | 26.29 |  | 17.07 |  |
| **S36** | Sputum | *H. influenzae*<br>(*H. influenzae*) | Undepleted | 22.58 | 9.70<br>(~10<sup>3</sup>) | 16.51 | 2.00 |
|  |  |  | Depleted | 32.28 |  | 18.51 |  |
| **S37** | Sputum | *P. aeruginosa*<br>(*P. aeruginosa*) | Undepleted | 21.56 | 11.69<br>(~10<sup>4</sup>) | 15.25 | 1.80 |
|  |  |  | Depleted | 33.24 |  | 13.45 |  |
| **S38** | Sputum | *S. aureus* & *P. aeruginosa*<br>(*S. aureus* & *P. aeruginosa*) | Undepleted | 20.76 | 6.87<br>(~10<sup>2</sup>) | 23.83 | 3.17 |
|  |  |  | Depleted | 27.63 |  | 20.66 |  |
| **S39** | Sputum | *H. influenzae*<br>(*H. influenzae* & *M. catarrhalis*) | Undepleted | 23.82 | 11.18<br>(~10<sup>3</sup>) | 14.45 | 2.79 |
|  |  |  | Depleted | 35.00 |  | 17.24 |  |
| **S40** | ETA | MRSA<br>MRSA | Undepleted | 21.69 | 4.28<br>(~19) | 19.91 | 1.62 |
|  |  |  | Depleted | 25.97 |  | 18.29 |  |
| **S41** | Sputum | *H. influenzae* & *S. aureus*<br>(*H. influenzae* & *S. aureus*) | Undepleted | 20.86 | 14.14<br>(~10<sup>4</sup>) | 16.71 | 6.85 |
|  |  |  | Depleted | 35.00 |  | 23.56 |  |

#### Reducing turnaround time

Prior to optimisation, the host DNA depletion method took approx. 1.5 hrs. Removal of the second DNase treatment and reducing the number of washes shortened this to 50 min without affecting the degree of human DNA depletion compared to the pilot method (average: >10^3^ fold;) (Supplementary Table 2a). Time could also be saved by reducing the library preparation PCR extension time from six to four minutes. Comparison of the microbial community profile after sequencing showed no difference in the top three most abundant species and only a small reduction in average fragment length (<400bp) between libraries produced with four and six minute extension times (Supplementary Table 2b). This enabled MinION library preparation to be completed within two hours and a half (reduced from 3.5 hours), giving an overall turnaround time of under four hours prior to DNA sequencing.

### Limit of detection

The limit of detection (LoD) of the method was determined using an uninfected ‘normal respiratory flora’ (NRF) sputum sample spiked with serial ten-fold dilutions of *S. aureus* and *E. coli* cultures at known cell densities. The LoD was defined as the fewest cells required for the identification of the infecting ‘pathogen’ using the analysis pipeline as follows: ≥1% microbial reads identified using WIMP and an alignment score ≥20. The LoD was determined to be 10,000 (10^4^) cells for both *E. coli* and *S. aureus* (Supplementary Table 3).

**Table 3:** Optimised metagenomic pipeline output compared to routine microbiology culture results

| Pathogen | Culture positive | Metagenomics |  |  |
| --- | --- | --- | --- | --- |
|  |  | Detected | Missed detection | Additional detection |
| *Haemophilus influenzae* | 10 | 10 | 0 | 4 (4*) |
| *Staphylococcus aureus* | 8 | 8 | 0 | 1 |
| *Pseudomonas aeruginosa* | 6 | 5 | 1 | 1 |
| *Escherichia coli* | 3 | 3 | 0 | 1 (1*) |
| *Moraxella catarrhalis* | 2 | 2 | 0 | 3 (1*) |
| *Serratia marscens* | 2 | 2 | 0 | 0 |
| *Klebsiella oxytoca* | 1 | 1 | 0 | 0 |
| *Klebsiella pneumoniae* | 1 | 1 | 0 | 1 |
| *Streptococcus pneumoniae* | 0 | 0 | 0 | 6 (4*) |
| None (NRF/NSG) | 0 | 0 | 0 | 10** |

\*Additional detection in NRF/NSG samples \*\*Two additional pathogens detected in three NRF/NSG samples

### Mock community detection

The Optimised method was tested in triplicate on a panel of common respiratory pathogens spiked into NRF sputum (~10^8^ cfu/ml per pathogen) to determine whether the saponin human DNA depletion method led to inadvertent loss of any bacterial DNA. We observed no bacterial DNA loss (average ΔCq <1) for any organisms (*E. coli, H. influenzae, K. pneumoniae, P. aeruginosa, S. aureus* and *S. maltophilia*) tested except *S. pneumoniae* where there was a 5.7-fold loss, (average ΔCq 2.52) between depleted and undepleted samples (Supplementary Table 4).

**Table 4:**
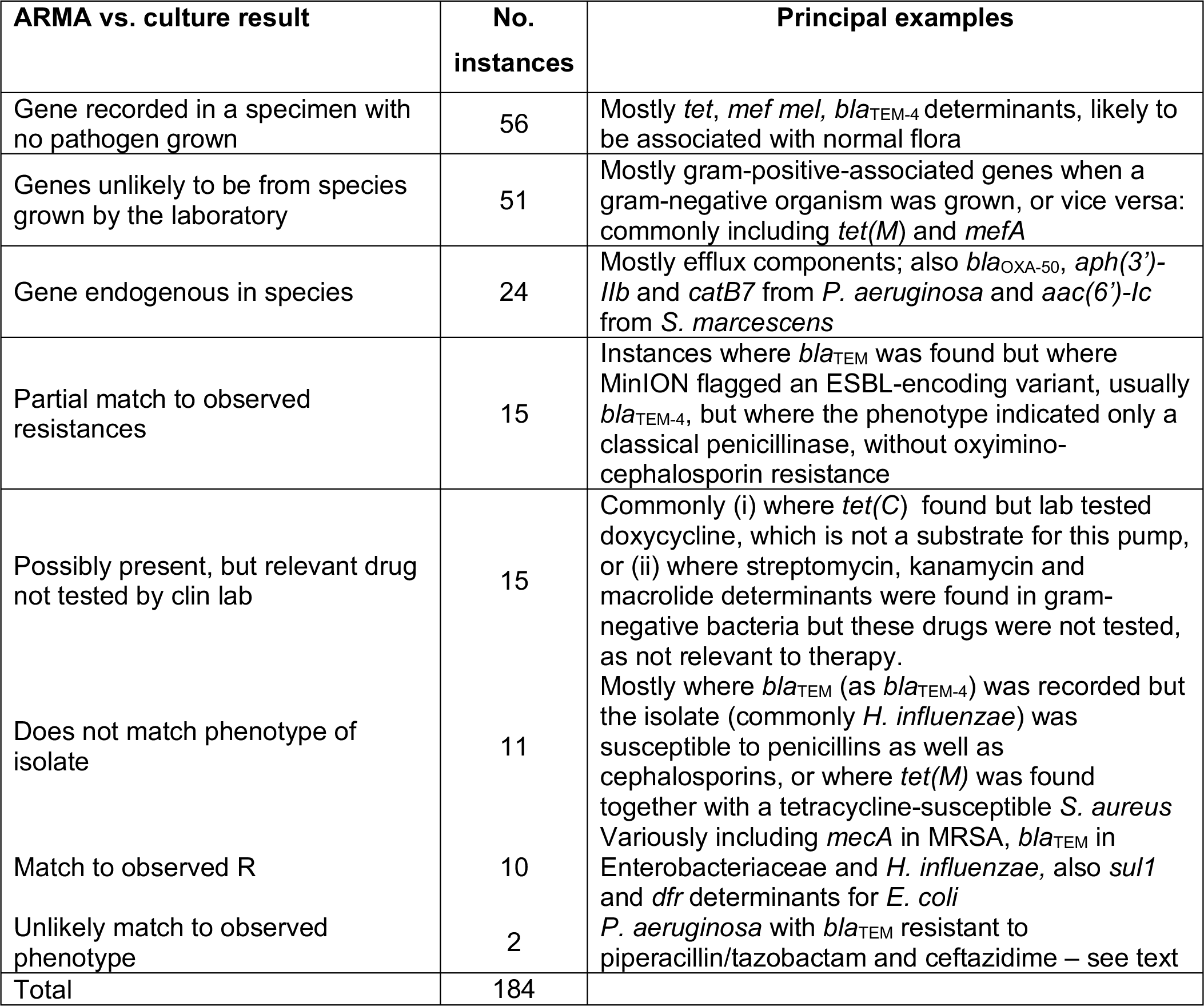
Resistance genes found by ARMA in relation to pathogens grown

### Optimised method evaluation

The optimised method was then tested on 41 respiratory samples from patients with suspected bacterial LRIs. A maximum of 10^4^ fold depletion of human DNA (average 10^3^ fold; range 4.8-18,054 fold) was observed between depleted and undepleted samples (Table 2). The overall sensitivity of the refined method for the detection of respiratory pathogens was 96.6% (95% CI, 80.4-99.8%), specificity was 41.7% (95% CI, 16.5-71.4%), again *not* counting additional organisms in culture-positive samples as false positives (Table 3). The turnaround time from sample to result was approx. 6 hours (including 2 hours MinION sequencing) (Supplementary Table 5).

The pathogenic organism reported by routine microbiology was detected together with an additional pathogen (not reported by culture) in eight samples: *K. pneumoniae* in S5, *P. aeruginosa* in S7, *M. catarrhalis* in S14 and S39, *S. aureus* in S29 and *S. pneumoniae* in S8 and S15 (Table 2). Up to two potentially pathogenic bacteria were also observed in seven samples reported as NRF/no significant growth (NSG) by routine microbiology, (S10, 11, 12, 21, 28, 31 and 32). *H. influenzae* and *S. pneumoniae* in S10 and S21; *S. pneumoniae* in S11 and S28; *M. catarrhalis* and *H. influenzae* in S12; *H. influenzae* in S31 and *E. coli* in S32. Only one pathogenic organism reported by routine microbiology was not detected using the Optimised method. Specifically, S9 was reported as a mixed infection with *P. aeruginosa* and *E. coli* whereas only *E. coli* was detected by metagenomics. There were two other mixed infections reported by routine microbiology, both involving *S. aureus* together with *H. influenzae* (S27 and S41), and both organisms were detected using the Optimised method.

### Antibiotic resistance analysis

The collection of samples tested using the optimised method had low rates of antibiotic resistance, as determined by routine testing (Supplementary table 6). Across the 33 cultivated organisms, just 38 instances of resistance or intermediate resistance to tested antibiotics were recorded (Table 4), and some of these overlapped (e.g. two *S. aureus* were resistant to oxacillin, flucloxacillin and penicillin).

Of these 38, five could be discounted as resistance inherent in the cultured species (e.g. to penicillins in *K. pneumoniae* and co-amoxiclav in *Serratia marcescens*). Of the remaining 33, 14 could be explained by the resistance genes found (all resistance genes were detected using ONT’s Antimicrobial Resistance Mapping Application – ARMA), including *mecA* found in both MRSA (S16 and S40), *sul1* and dfrA12 or *dfrA17* genes found in each of two co-trimoxazole resistant *E. coli* (S1 and S9), *aac(3’)-IIa* (and *IIc*) in a tobramycin-resistant *E. coli* (S9) and *bla*_TEM_ variants found in each of two amoxicillin-resistant *E. coli* (S1 and S35) and amoxicillin-resistant *H. influenzae* (S18 and S36). A caveat apropos *bla*_TEM_ is that ARMA mostly flagged *bla*_TEM-4_ when *bla*_TEM-1_ was considerably more likely given its vastly greater prevalence and the fact that the isolates remained susceptible to oxyimino-cephalosporins, which are substrates for TEM-4. Another 6/33 cases concerned non-susceptibility to penicillin/β-lactamase inhibitor combinations in isolates where *bla*_TEM_ or (*K. oxytoca*) *bla*_OXY_ was found; resistance here may be associated with these enzymes, but depends on their level of expression, which was not investigated (or measurable) by the present pipeline. Next, 2/33 instances concerned a specimen where *P. aeruginosa* (S37) was cultured and found resistant to ceftazidime and piperacillin/tazobactam: *bla*_TEM-4_ was found and could explain this phenotype but is unlikely in *P. aeruginosa*, where β-lactam resistance mostly reflects up-regulation of chromosomal *ampC* or efflux.

This leaves 11/33 instances where observed resistance was not explained by the genes found. Two of these were amoxicillin-resistant *M. catarrhalis* (S8 and S26), where genes for the likely BRO β-lactamases are absent from the ARMA databases. Another six, variously including ampicillin and co-trimoxazole resistance in *H. influenzae* (S7, S18, S36, S39 and S41) and resistances to trimethoprim, ciprofloxacin and fusidic acid in *S. aureus* (S16) were phenotypes where modification of chromosomal genes is the likely source of resistance, and the relevant sequence data is not analysed by ARMA. Last, there were three instances - one *S. aureus* resistant to gentamicin (S16) and a *K. pneumoniae* (S2) resistant to both co-amoxiclav and piperacillin/tazobactam but lacking any acquired β-lactamase, where we cannot explain the failure of sequencing to identify a likely mechanism.

Looked at another way, sequencing identified 184 resistance genes across the 41 specimens, based on multiple inclusion when ARMA identified multiple variants of e.g. *bla*_TEM_ in a specimen. Many of these genes are likely to have originated from the normal flora, as evidenced by the fact that *tet(M)* was found in 8/12 NRF/NSG specimens as was *bla*_TEM-4_, whilst *mefA* and *mel* were each found in 9/12 such samples. Other genes, including sul1, dfr determinants and *mecA* were, however, only found in association with cultivation of an organism with the corresponding resistance. Endogenous resistance genes of cultivated species were widely seen, such as *bla*_OXA-50_, *catB7* and *aph(3’)-IIb* in *P. aeruginosa* (S3, S29, S30 and S37) and *aac(6’)-Ic* in *S. marcescens* (S4 and S13).

The specificity and sensitivity of the developed method for resistance gene detection was not determined as this would have required isolating and sequencing all bacteria (pathogens and commensals) present in the sputum samples, which was beyond the scope of the study.

### Reference-based genome assembly

Two samples containing antibiotic resistant bacteria were chosen as examples to generate reference-based genome assemblies directly from the metagenomic data. Assemblies were generated for an MRSA (S16) and an *E. coli* resistant to amoxicillin, co-amoxiclav and co-trimoxazole (S1). The results were compared with those for undepleted controls after two and 48 hours of sequencing. Within the first two hours of sequencing the human DNA depleted MRSA sample had 64.7x genome coverage with an assembly of 64 contigs (longest contig = 1360276 and N50=106kbp, data not shown). Genome coverage increased to 232.5x after 48hrs of sequencing, with a final assembly consisting of 22 contigs (longest contig = 481kbp and N50=402kbp). After 48 hrs of sequencing both the depleted and undepleted MRSA samples had sufficient coverage to enable assembly, though coverage was much greater for the depleted sample (232.5x) than the undepleted (18.5x) with a final assembly of 34 contigs (longest contig = 416kbp and N50=178kbp) (Figure 2a) At 2hrs of sequencing the undepleted MRSA sampe had an assembly of 216 contigs (longest contig = 32kbp and N50=7kbp, data not shown).

**Figure 2:**
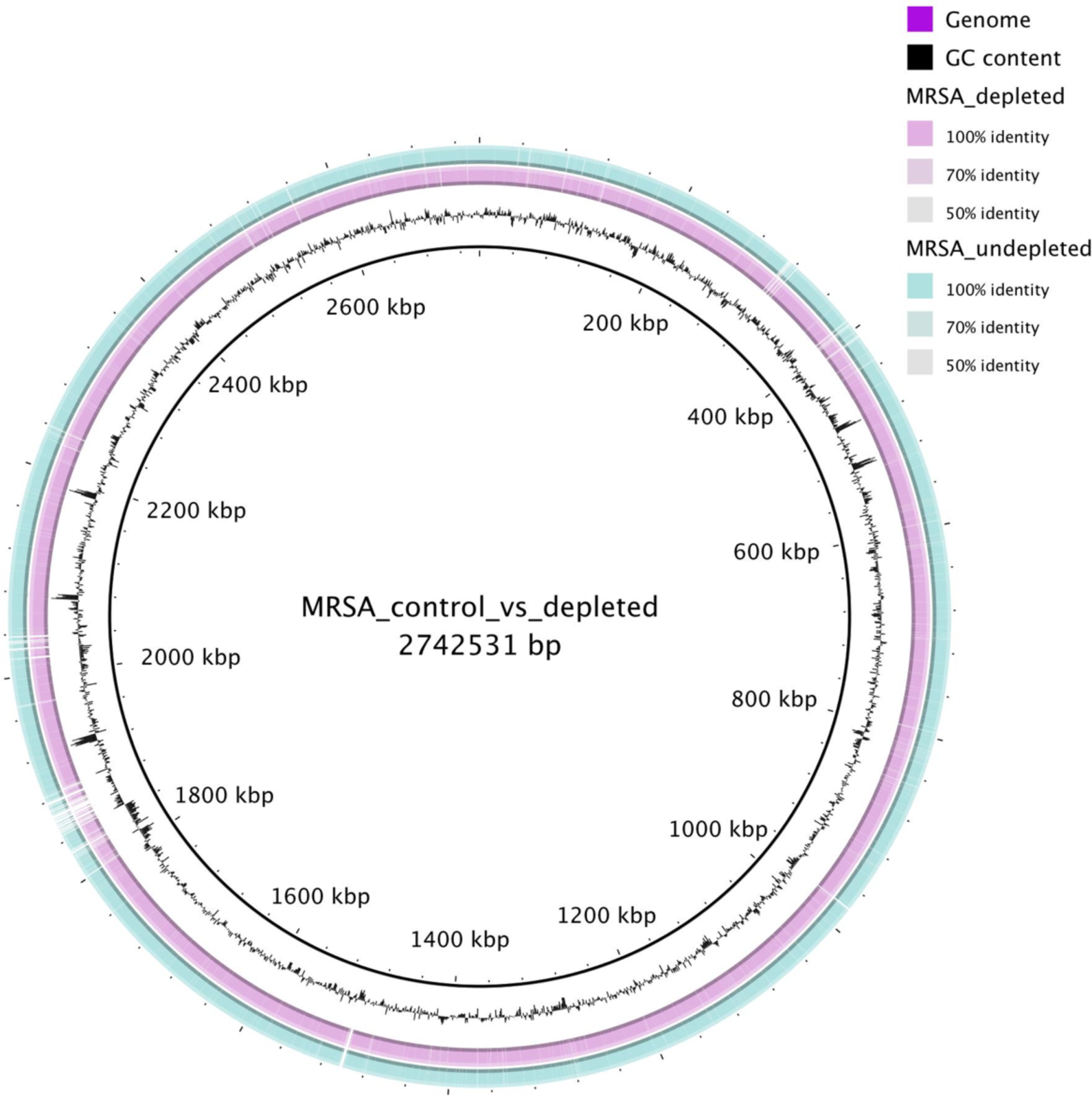
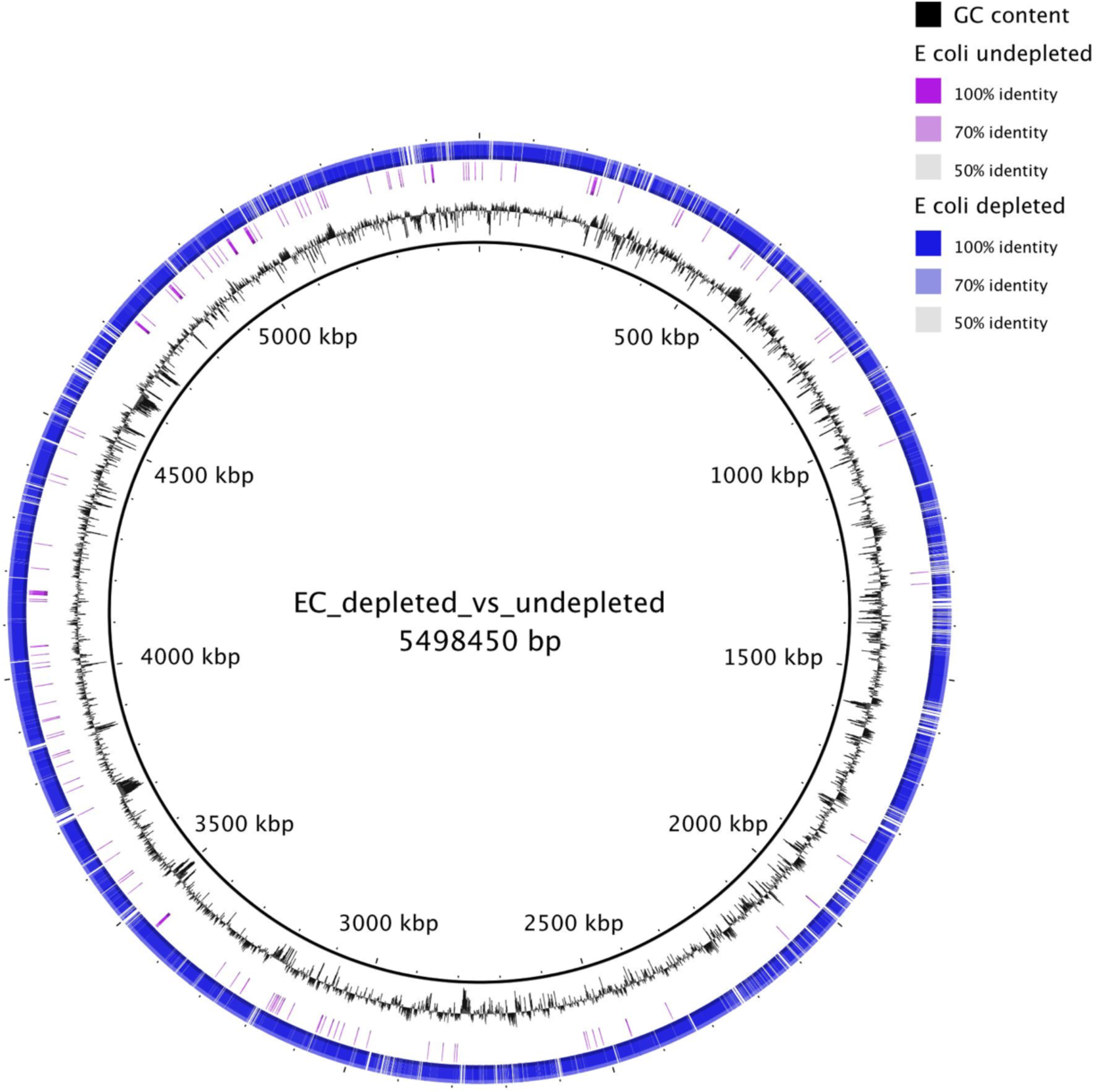
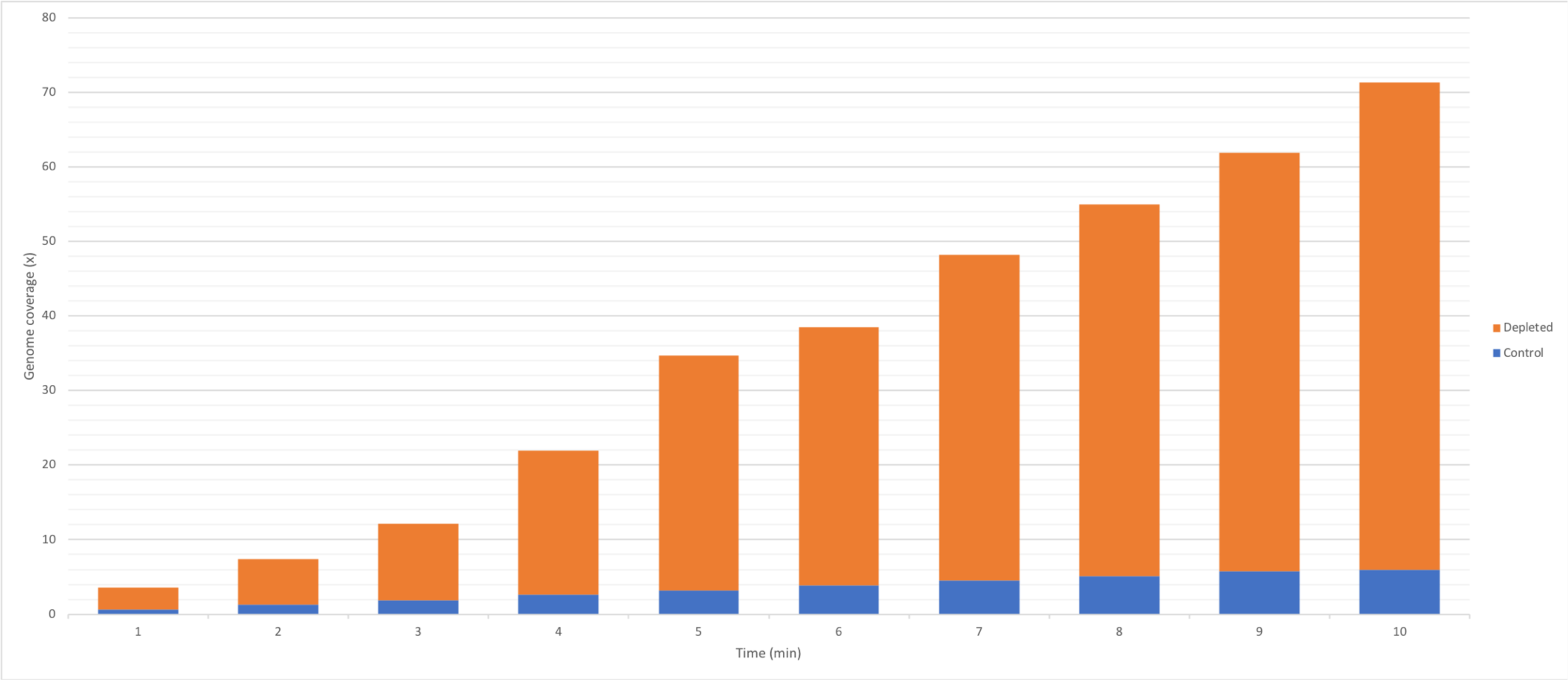
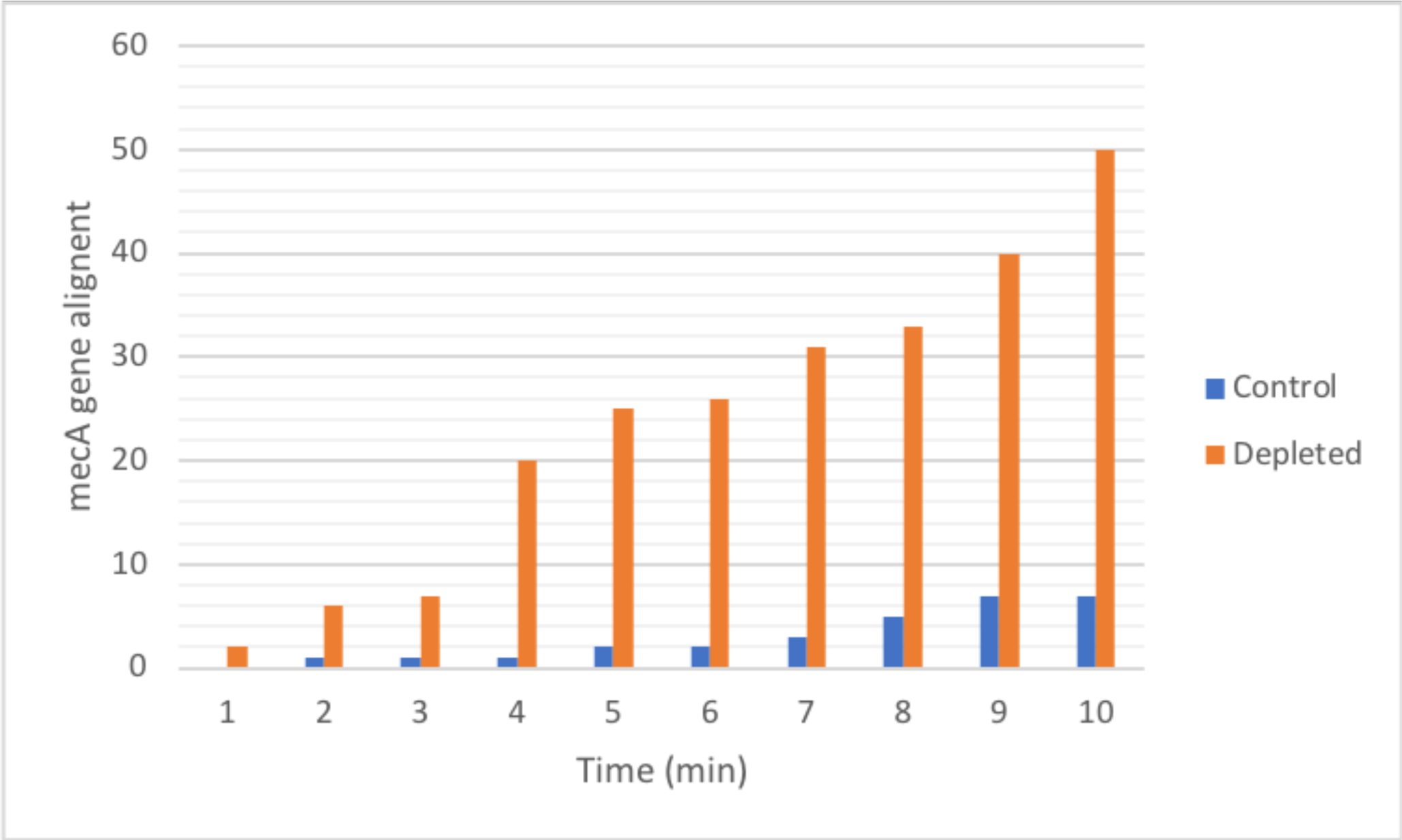
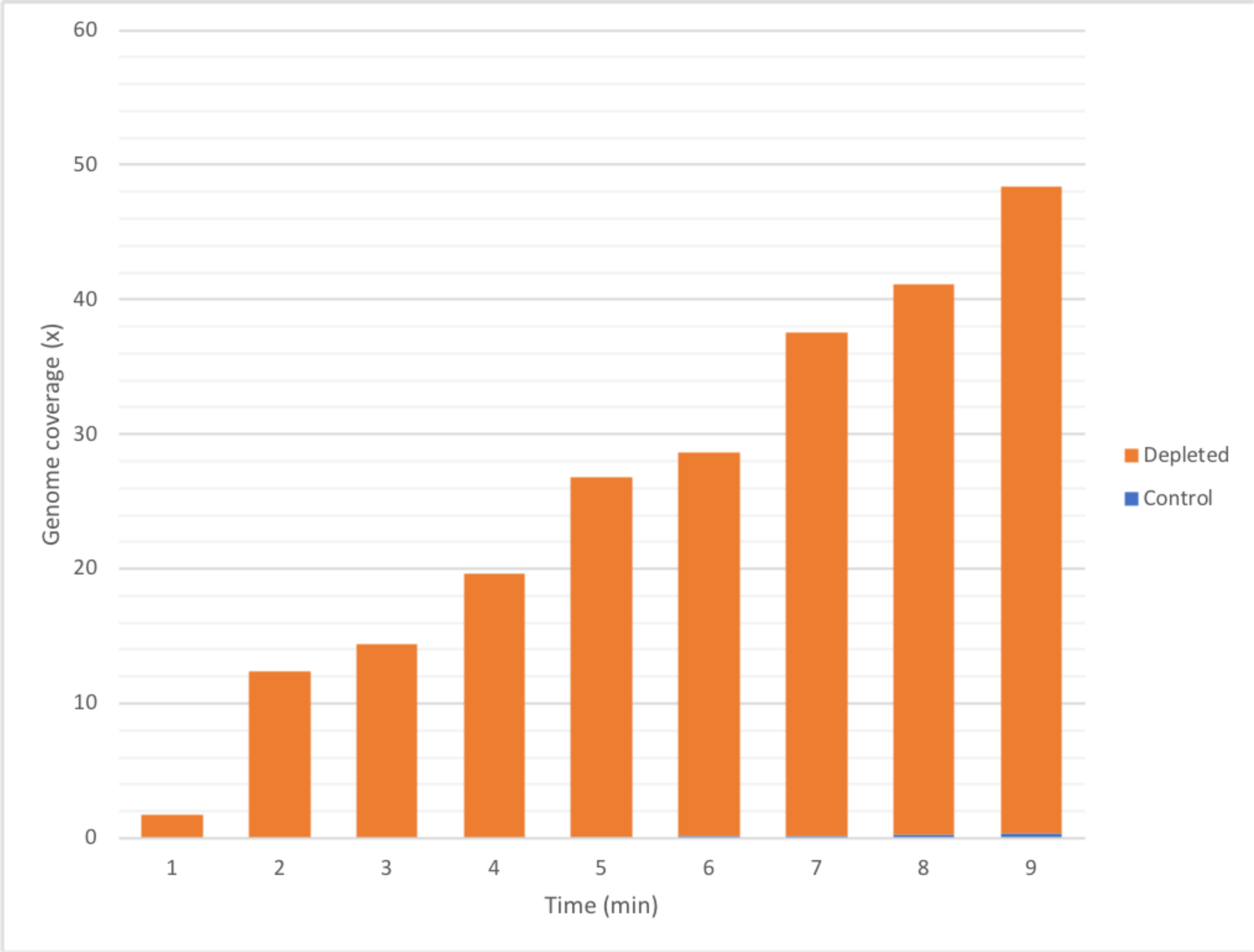
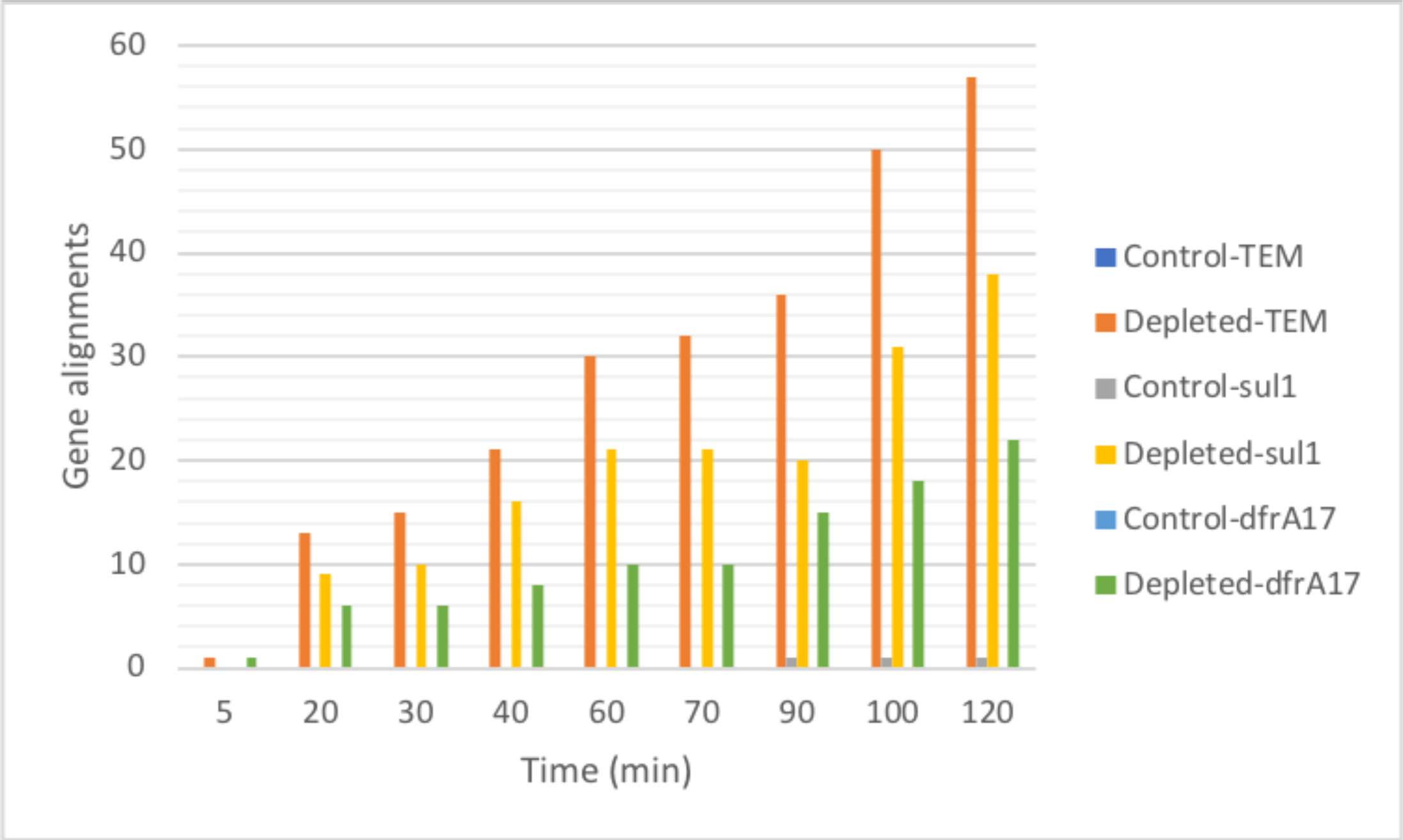
Bacterial genome assembly, genome coverage and antibiotic gene detection with depleted versus undepleted samples. A: MRSA after 48 hours of sequencing. B: *E. coli* after 48 hours of sequencing. C: MRSA genome coverage of depleted versus undepleted during two hours of sequencing. D: *mecA* gene alignment of depleted versus undepleted during two hours of sequencing. E: *E. coli* genome coverage of depleted versus undepleted during two hours of sequencing. F: *bla*_TEM_, *sul1* and *dfr*A17 gene alignment of depleted versuss undepleted during two hours of sequencing.

For the sample positive for resistant *E. coli* there was 21x genome coverage within two hours, with an assembly of 83 contigs (longest contig = 437kbp and N50=164kbp, data not shown). Genome coverage increased to 165x after 48 hrs with the final *E. coli* assembly having 78 contigs (longest contig = 473kbp and N50=177kbp). The undepleted sample data only produced 0.3x coverage after 2hrs, which increased to 1.2x genome coverage after 48 hrs sequencing (Figure 2b).

### Time-point analysis

Using the same sample set as for genome assembly, data from the first two hours of sequencing were compared over time for depleted samples and undepleted controls. Within 5 min of sequencing the depleted MRSA sample (S16) had 3x genome coverage compared with the undepleted control (S16-undepleted) at 0.6x coverage (Figure 2c). The *mec*A gene was not detected in the undepleted sample after 5 min whereas two *mec*A gene alignments were detected in the depleted sample by the same time point (Figure 2d).

The depleted *E. coli* sample (S1) had 1.74x genome coverage within 5 min of sequencing whereas the undepleted control (S1-undepleted) had 0.03x (Figure 2e). This *E. coli* was resistant to amoxicillin (*bla*_TEM_ gene), co-amoxiclav (possibly owing to *bla*_TEM_ if strongly expressed) and co-trimoxazole (*sul1* and *dfr*A17 genes). The *bla*_TEM_ and *dfr*A17 genes were not detected in the undepleted sample within two hours of sequencing and only one alignment was detected for *sul*1. Conversely, all three resistance genes were detected within 20 min of sequencing in the depleted sample and, after two hours, 57 *bla*_TEM_, 38 *sulf1* and 22 *dfrA17* alignments were detected (Figure 2f).

### Cost

The running cost of the optimised pipeline developed in this study was approximately $130 per sample when run in batches of six per flowcell ($546 for singleplex runs) (Supplementary Table 7).

## Discussion

Current culture-based diagnostics and susceptibility testing, though used for 70 years ^29^, have serious limitations as guides for the appropriate clinical management of acute serious infection. This is mainly due to slow sample-to-result turnaround. Rapid, accurate diagnostics are urgently required, so that patients can be prescribed appropriate antibiotics as quickly as possible, potentially improving both outcomes and antimicrobial stewardship. We developed and tested a novel nanopore sequencing-based clinical metagenomics pipeline for the microbiological investigation (pathogen and antibiotic resistance gene identification) of bacterial LRIs within 6h of clinical diagnosis (assuming the sample is tested immediately).

We began by developing an efficient method for the removal of host DNA from sputum, bronchoalveolar lavage (BAL) and endotracheal aspirates (ETAs). This saponin-based differential lysis method resulted in ~10^3^-fold host DNA depletion without any significant bacterial loss. This high efficiency depends upon the high concentrations of saponin and nuclease buffer salt used. A very high salt buffer (5.5M NaCl) was found to be optimal for nuclease (HL-SAN) digestion of DNA, even though this was far beyond the manufacturer’s recommended range for optimum activity. This, combined with a final saponin concentration of 2.2-2.5%, which is higher than used in other saponin based depletion methods ^19^, resulted in efficient leukocyte lysis and efficient human DNA depletion. It should be noted that, the same method depletes approx. 10^5^-fold human DNA on average from less inhibitory sample types such as blood and tissue (data not shown). Forty LRI samples were sequenced using the initial Pilot method, which was 91.2% sensitive and 83.3% specific for the detection of respiratory pathogens compared with “gold standard” culture.

The Pilot method was then optimised, by shortening the depletion protocol, introducing bead-beating for improved organism lysis, and reducing the library preparation time by shortening the PCR extension time. The Optimised method was 96.6% sensitive with a turnaround time of six hours. Only one pathogen (1/41) detected by culture was missed (false-negative major error) using the Optimised method (vs. 3/40 using Pilot method); specifically, a *P. aeruginosa*, which was found by culture in a mixed infection with *E. coli*. Possible explanations for missing this pathogen are (i) PCR competition/bias leading to preferential amplification of DNA from the more abundant pathogen or (ii) loss of pseudomonal DNA during host depletion, caused by damaged but viable cells sensitive to saponin lysis. Nevertheless, the mock community experiments demonstrated that common respiratory pathogens including *P. aeruginosa* were not effected by saponin depletion, the sole notable exception being *S. pneumoniae*. Under stress, *S. pneumoniae* is prone to autolysis through the activation of a pneumolysin gene, and this may occur to a portion of the bacteria during the host DNA depletion process ^30^ (but may also have been caused by autolysis of *S. pneumoniae* when growing to stationary phase for mock community experiments). *S. pneumoniae* was correctly identified in five of six patients found by culture to be infected with *S. pneumoniae*, but it may have been underrepresented in these samples. The time from sample collection to bacterial DNA extraction may be a critical factor for the accurate detection of *S. pneumoniae*.

The specificity of the Optimised method was significantly lower than the Pilot method (41.6% vs 83.3%). This is likely to be related to the increased sensitivity of the Optimised method (96.5% vs 91.2%). Metagenomic detection of additional pathogens compared to culture was expected as the clinical laboratory routinely dilutes the sample (1/1000) before plating and we used a larger sample volume (400µl vs 10µl). Most (10/17) of the additional organisms detected were potential known pathogens that can also be carried as commensals in the respiratory tract, notably including *S. pneumoniae* and *H. influenzae* ^31–33^. Hence, it is likely that the additional pathogens not detected by culture were present in the samples and that culture either failed to detect them or did not report them as significant. However, some of the additional detection observed might be due to bioinformatic misclassification, as *k-mer* based read classification can be unreliable at the species level, particularly where species in a genus are highly related or share genes. This is particularly important in samples containing high numbers of commensal Streptococci/Haemophilus where a proportion of commensal reads can be misclassified as pathogen reads ^34, 35^.

Cut-offs, in terms of number of bacteria per ml of body fluid, are applied in clinical microbiology laboratories for some infections including those of the urinary and respiratory tracts. The same approach is required for metagenomics. The clinical cut-off used for respiratory samples is typically 10^4^ pathogens per ml of sample (range 10^3^-10^5^ per ml dependent on sample type – achieved by sample dilution) ^36^. We routinely applied cut-offs at 1% of classified reads, with a WIMP alignment score ≥20. We chose these cut-offs to: censor reads arising from pipeline contaminants; remove barcode leakage between samples on multiplexed runs (ONT’s Flongle (https://nanoporetech.com/products/comparison), an adapter for single use flowcells designed for diagnostic applications, should overcome this issue) and; remove low quality WIMP alignments, which result in misclassified reads. Results from our LoD experiments for the Optimised pipeline (10^4^ cells/ml) are within the range of culture-based clinical cut-off. More precise methods for identifying pathogen species in clinical metagenomic data are urgently required.

To maximise the impact on patient management, identification of clinically relevant antibiotic resistance genes as well as the infecting pathogen/s is necessary. In this regard the present pipeline has potential but requires refinement. Both MRSA cases were identified by the presence of *mecA*, with no false positives for this gene. Co-trimoxazole resistance in Enterobacteriaceae was accurately identified with detection of *sul* and *dfr* genes and these were not found in *H. influenzae*, for which resistance is largely mutational ^37, 38^. However, genes such as *tet(M), mel, mefA* and *bla*_TEM_ were found in 8/12 samples where no pathogen was grown, suggesting presence in the normal or colonising respiratory flora. To overcome this issue, it would be necessary to associate resistance genes to particular organisms. This can be done by examining flanking sequences ^39–42^ in the c. 3 kb nanopore reads in cases where a gene is chromosomally inserted, as is usual for transposon borne *tet(M)* and *mefA* in streptococci ^43–45^ (Supplementary Figure 1), including *S. pneumoniae*, but will not determine the source of plasmid-borne resistance genes. Real-time consessus calling for the detection of mutational resistance and chromosomal resistance gene source determination functionality would significantly improve the accuracy of the ARMA output.

An additional advantage of implementing clinical metagenomics (compared to other rapid tests e.g. PCR) is the impact it could have beyond clinical diagnosis. To illustrate this potential, the data generated after 48hrs of sequencing was used to perform reference based pathogen genome assemblies, identifying bacteria beyond species level ^17^. Such information is equivalent to reference laboratory typing which is, in the UK, carried out by whole genome sequencing of isolates. The quality of the metagenomic data generated by the method reported here would allow investigations of the emergence and patient-to-patient spread of pathogens and antimicrobial resistance directly from clinical samples in real-time ^46, 47^. Currently it would be necessary to run “reference” culture alongside PCR based approaches, otherwise the link to phenotypes would be lost – clinical metagenomics, on the other hand, has the potential to replace routine culture, allowing it to be reserved for more unusual pathogens/resistances.

In conclusion, we report the first rapid clinical metagenomics pipeline for the microbiological investigation of bacterial LRIs. The method provides accurate pathogen and antibiotic resistance gene identification within six hours. With additional sequencing time (up to 48 hrs), it provides sufficient data for public health and infection control applications. The pipeline is currently being evaluated in a clinical trial (INHALE - http://www.ucl.ac.uk/news/news-articles/1115/181115-molecular-diagnosis-pneumonia) on the rapid diagnosis of hospital-acquired and ventilator-associated pneumonia.

## Methods

### Ethics

This study used excess respiratory samples, after routine microbiology diagnostic tests had been performed, from patients with suspected LRIs such as persistent (productive) cough, bronchiectasis, CAP/HAP, cystic fibrosis and exacerbation of chronic obstructive pulmonary disease (COPD, emphysema/chronic bronchitis). Ethical approval was not required as the samples were used for method development, and no patient identifiable information was collected. The only data collected were routine microbiology results, which detailed the pathogen(s) identified and their antibiotic susceptibility profiles.

### Routine clinical microbiological investigation

Respiratory samples including sputum, endotracheal secretions and ETAs were treated with sputasol (Oxoid-SR0233) in a 1:1 ratio before being incubated for a minimum of 15 min at 37°C. Sputasol-treated respiratory samples (10 µl) were inoculated into 5 ml of sterile water and mixed. Following this, 10 µl of sample was streaked onto blood, chocolate and cysteine lactose electrolyte deficient (CLED) agar. BAL samples, they were not treated with sputasol; instead they were centrifuged to concentrate bacterial cells for a minimum of 10 min at 3000 rpm. BALs did not undergo further dilution and were streaked directly onto the agar plate. Depending on clinical details and the source of the specimen, other agar plates (including sabouraud, mannitol salt and *Burkholderia cepacia* selective agar) were additionally used.

All inoculated agar plates were incubated at 37 ̊C overnight and then examined for growth with the potential for re-incubation up to 48 hours. If any significant organism was grown then antibiotic susceptibility testing by agar diffusion using EUCAST methodology was performed. The laboratory’s Standard Operating Procedure is based on the Public Health England UK Standards for Microbiology Investigations B 57: Investigation of bronchoalveolar lavage, sputum and associated specimens ^36^.

### Sample collection and storage

The excess respiratory samples (sputa, ETA, BAL) were collected after culture and susceptibility testing at Norfolk and Norwich University Hospitals (NNUH) Microbiology Department (described above) and stored at 4 °C prior to testing. They were indicated by clinical microbiology to contain bacterial pathogen(s), NRF or to have yielded NSG. Forty samples (n=34 positive and n=6 NRF samples, comprising 34 sputa, four BALs and two ETAs) were used to test the first ‘Pilot” iteration of the diagnostic pipeline and another 41 (n=29 suspected LRI, n=9 NRF and n=3 NSG samples, comprising 38 sputa, one BAL and two ETAs) to test the Optimised pipeline.

### First iteration of the diagnostic pipeline (Pilot method): Host DNA Depletion

Respiratory samples (400 µl) were centrifuged at 8000 xg for 5 min, after which the supernatant was carefully removed and the pellet resuspended in 250 µl of PBS, any pellet >50 µl was diluted one in four and re-centrifuged. The saponin-based differential lysis method was modified from previously reported saponin methods ^48, 49^: saponin (Tokyo Chemical Industry-S0019) was added to a final concentration of 2.5 % (200 µl of 5 % saponin), mixed well and incubated at room temperature (RT) for 10 min to promote host cell lysis. Following this incubation, 350 µl of water was added and incubation was continued at RT for 30 s, after which 12 µl of 5 M NaCl was added to deliver an osmotic shock, lysing the damaged host cells. Samples were next centrifuged at 6000 xg for 5 min, with the supernatant removed and the pellet resuspended in 100 µl of PBS. HL-SAN buffer (5.5 M NaCl and 100 mM MgCl_2_ in nuclease-free water) was added (100 µl) with 5 µl HL-SAN DNase (25,000 units, Articzymes - 70910-202) and incubated for 15 min at 37°C with shaking at 800 RPM for host DNA digestion. An additional 2 µl of HL-SAN DNase was added to the sample, which next was incubated for a further 15 min at 37°C with shaking at 800 RPM. Finally, the host-DNA depleted samples were washed three times with decreasing volumes of PBS (300 µl, 150 µl, 50 µl). After each wash, the sample was centrifuged at 6000 xg for 3 min, the supernatant discarded and the pellet resuspended in PBS.

### Pilot method: Bacterial Lysis and DNA Extraction

After the final wash step of the host depletion, the pellet was resuspended in 380 µl of bacterial lysis buffer (Roche UK-4659180001) and 20 µl of proteinase K (>600mAu/ml) (Qiagen −19133) was added before incubation at 65°C for 10 min with shaking at 800 RPM (on an Eppendorf Thermomixer). Nucleic acid was then extracted from samples using the Roche MagnaPure Compact DNA_bacteria_V3_2 protocol (MagNA pure compact NA isolation kit I, Roche UK-03730964001) on a MagNA Pure Compact machine (Roche UK-03731146001).

### Optimised method: Host DNA Depletion (Figure 1)

The optimized method sought to improve and shorten some steps. Specifically, after the first 5 min centrifugation at 8000 x g, up to 50 µl of supernatent was left so as to not disturb the pellet (final saponin conc. 2.2-2.5%). Instead of performing two rounds of host DNA digestion, the amount of HL-SAN DNase was increased up to 10 µl and a single incubation of 15 min at 37 °C was carried out with shaking at 800 RPM on an Eppendorf Thermomixer. Finally, the number of washes was reduced to two with increasing volumes of PBS (800 µl and 1 ml).

### Optimised method: Bacterial Lysis and DNA Extraction (Figure 1)

After the final wash, the pellet was re-suspended in 500 µl of bacterial lysis buffer (Roche UK - 4659180001), transferred to a bead-beating tube (Lysis Matrix E, MP Biomedicals - 116914050) and bead-beaten at maximum speed (50 oscillations per second) for 3 min in a Tissue Lyser bead-beater (Qiagen - 69980). This ensured the release of DNA from difficult-to-lyse organisms (e.g. *S. aureus*). The sample was centrifuged at 20,000 xg for 1 min and ~230 µl of supernatant was transferred to a fresh Eppendorf tube. The volume was topped-up with 170 µl of bacterial lysis buffer and 20 µl of proteinase K (>600 mAu/ml, Qiagen - 19133) was added. Samples were then incubated at 65°C for 5 min with shaking at 800 RPM on an Eppendorf Thermomixer. DNA was extracted from samples using the Roche MagnaPure Compact DNA_bacteria_V3_2 protocol (MagNA pure compact NA isolation kit I, Roche UK - 03730964001) on a MagNA Pure Compact machine (Roche UK −03731146001).

### DNA quantification and quality control

DNA quantification was performed using the high sensitivity dsDNA assay kit (Thermo Fisher - Q32851) on the Qubit 3.0 Fluorometer (Thermo Fisher - Q33226). DNA quality and fragment size (PCR products and MinION libraries) were assessed using the TapeStation 2200 (Agilent Technologies - G2964AA) automated electrophoresis platform with the Genomic ScreenTape (Agilent Technologies - 5067-5365) and a DNA ladder (200 to >60,000 bp, Agilent Technologies - 5067-5366).

### MinION Library Preparation and Sequencing

MinION library preparation was performed according to the manufacturer’s instructions for (i) the Rapid Low-Input by PCR Sequencing Kit (SQK-RLI001), (ii) the Rapid Low-Input Barcoding Kit (SQK-RLB001) or (iii) the Rapid PCR Barcoding Kit (SQK-RPB004) with minor alterations as follows. For single sample sequencing runs using the SQK-RLI001 kit, 10 ng of the MagNA Pure-extracted DNA were used for the tagmentation/fragmentation reaction, where DNA was incubated at 30°C for 1 min and at 75°C for 1 min. The PCR reaction was run as per the manufacturer’s instructions; however, the number of PCR cycles was increased to 20. For multiplexed runs, SQK-RLB001 and SQK-RPB004 kits were used. A 1.2x AMPure XP bead (Beckman Coulter-A63881) wash was introduced after the MagNA Pure DNA extraction and prior to library preparation for multiplexed runs and DNA was eluted in 15 µl of nuclease-free water. Modifications for the library preparation were i) 10 ng of input DNA and 2.5 µl of FRM were used for the tagmentation/fragmentation reaction and nuclease-free water was used to make the volume up to 10 µl, ii) for the PCR reaction, 25 cycles were used and the reaction volume was doubled. All samples run using the Pilot method used a 6 min extension time, whereas the Optimised method used a reduced extension time of 4 min. When multiplexing, PCR products were pooled together in equal concentrations, then subjected to a 0.6x AMPure XP bead wash and eluted in 14 µl of the buffer recommended in the manufacturer’s instructions (10 μL 50 mM NaCl, 10 mM Tris.HCl pH8.0). Sequencing was performed on the MinION platform using R9.4, R9.5 or R9.4.1 flow cells. The library (50-300 fmol) was loaded onto the flow cell according to the manufacturer’s instructions. ONT MinKNOW software (versions 1.4-1.13.1) was used to collect raw sequencing data and ONT Albacore (versions 1.2.2-2.1.10) was used for local base-calling of the raw data after sequencing runs were completed. The MinION was run for up to 48 hours with WIMP/ARMA analysis performed on the first six folders (~24,000 reads) for Pilot method samples and the first two hours of data for all Optimised method samples.

### Quantitative PCR (qPCR) assays

Probe or SYBR Green based qPCR was performed on samples to detect and quantify human DNA, DNA targets for specific pathogens (*E. coli*, *H. influenzae, K. pneumoniae, P. aeruginosa, S. aureus, Stenotrophomonas maltophilia* and *S. pneumoniae*) and the bacterial 16S rRNA V3-V4 gene fragment. All qPCR assays were performed on a Light Cycler^®^ 480 Instrument (Roche). Details of primer sequences and targets can be found in Supplementary Table 8 (oligonucleotides were supplied by Sigma or ThermoFisher Scientific).

For all probe-based qPCR reactions, the master mix consisted of 10 µl LightCycler 480 probe master (2X), 0.5 µl each of reverse and forward primer (final conc. 0.25 µM) and 0.4 µl probe (final conc. 0.2 µM). For all SYBR-Green-based qPCR reactions, the master mix consisted of 10 µl LightCycler 480 SYBR Green I master (2x) and 1 µl of each forward and reverse primer (final conc. 0.5 µM). To the PCR mix, 2 µl of DNA template and nuclease-free water to a total volume of 20 µl were added. The qPCR conditions were: pre-incubation at 95°C for 5 min, amplification for 40 cycles at 95°C for 30 sec, 55°C for 30 sec and 72°C for 30 sec, with a final extension at 72°C for 5 min. Melt curves analysis was performed at 95°C for 5 sec, 65°C for 1 min, ramping to 95°C at 0.03°C/s in continuous acquisition mode, followed by cooling to 37°C.

### Example Limit of detection

The limit of detection (LoD) of the Optimised method for the detection of one Gram-positive and one Gram-negatie bacteria in sputum was determined using serial dilutions (10 –10^5^ cfu/ml) of cultured *E. coli* (H141480453) and *S. aureus* (NCTC 6571). Serial dilutions were made in sterile PBS and plated in triplicate on LB agar to determine colony forming units (CFU) per ml. The same dilutions were used to spike an NRF sputum sample for LoD experiments. Detection and quantification of bacterial DNA was performed using probe-based qPCR assays and MinION sequencing.

### Mock community experiments

Clinical isolates from respiratory samples were used to generate a mock community consisting of *S. pneumoniae, K. pneumoniae, H. influenzae, S. maltophilia* and *P. aeruginosa. E. coli* and *S. aureus* strains were also included (H141480453 and NCTC 6571 respectively). Pathogens (*E. coli* and *S. aureus* in 10 ml Luria-Broth and *K. pneumoniae*, *P. aeruginosa* and *S. maltophilia* in 10 ml Tryptic Soy Broth (TSB)) were cultured overnight at 37°C with shaking at 180 RPM. *H. influenzae* (in 10 ml TSB) and *S. pneumoniae* (in 10 ml Brain Heart Infusion Broth) were cultured statically at 37°C with 5% CO_2_ in an aerobic incubator. Cultured pathogens were then spiked into an NRF sample (~10^8^ cfu/ml per pathogen). The spiked samples were then tested in triplicate with the Optimised method, to determine if saponin depletion resulted in any inadvertent lysis of pathogens and loss of their DNA. All spiked samples were processed alongside undepleted controls. Probe or SYBR Green-based qPCR assays were used to determine the relative quantity of each spiked pathogen in depleted and undepleted spiked sputum samples.

### Pathogen identification and antibiotic resistance gene detection

The EPI2ME Antimicrobial Resistance pipeline (ONT, versions 2.47.537208-2.52.1202033) was used for initial analysis of MinION data for the identification of bacteria present in the sample and any associated antimicrobial resistance genes. Within this pipeline, WIMP (What’s in my Pot – which utilises ‘Centrifuge’ *kmer*-based read identification ^50^) was used for respiratory pathogen identification and ARMA (Antimicrobial Resistance Mapping Application –which utilised the CARD database ^51^) for antibiotic resistance gene detection. Potential pathogen(s) were reported if the number of reads accounted for ≥1% of microbial reads and with a WIMP alignment score ≥20. Antibiotic resistance genes were reported if >1 gene alignment was present using the ‘clinically relevant’ parameter within ARMA. This currently only reports resistance genes, acquired and chromosomal, but does not report resistance mutations/SNPs.

### Bacterial genome assembly

Genome assembly was performed first using Fast5-to-Fastq to remove reads shorter than 2000 bp and with a mean quality score lower than seven (https://github.com/rrwick/Fast5-to-Fastq). Porechop was used to remove sequencing adapters in the middle or the ends of each read, and re-identification of barcodes was carried out for each multiplexed sample (v0.2.3) (https://github.com/rrwick/Porechop). Filtered reads were aligned to a reference genome (chosen based on WIMP classification of pathogen reads) using Minimap2 with default parameters for ONT long-read data (v2.6-2.10) ^52^. Finally, Canu was used to assemble mapped reads into contigs using this long-read sequence correction and assembly tool (v1.6) ^53, 54^. BLAST Ring Image Generator (BRIG) was used for BLAST comparisons of the genome assemblies generated ^55^.

### Human DNA removal

Human DNA reads were removed from basecalled FASTQ files using Minimap2 to align to the human hg38 genome (GCA_000001405.15 “soft-masked” assembly) prior to Epi2ME analysis. Only unassigned reads were exported to a bam file using Samtools (-f 4 parameter). Non-human reads were converted back to FASTQ format using bam2fastx. These FASTQ files were processed for pathogen identification using WIMP and antibiotic resistance gene detection with ARMA. Further downstream analysis for genome coverage was performed using Minimap2 with default parameters for long-read data (-a -x map-ont) and visualised using qualimap. All clinical sample sequence data and assemblies are available at figshare with human DNA reads removed:

pilot method data: https://doi.org/10.6084/m9.figshare.6825410.v1

limit of detection: https://doi.org/10.6084/m9.figshare.6825389.v1

refined method data: https://doi.org/10.6084/m9.figshare.6825470.v1

bacterial genome assemblies: https://doi.org/10.6084/m9.figshare.6825323.v1).

## Acknowledgements

This study was supported by the UK Antimicrobial Resistance Cross Council Initiative (JOG, GK - MR/N013956/1), Rosetrees Trust (JOG - A749), NIHR INHALE study (JOG, HR, DL, RB - RP-PG-0514-20018), the University of East Anglia (JOG, TC) and Oxford Nanopore Technologies (JOG, TC, DT). Part of the bioinformatics analysis was run on CLIMB-computing servers, an infrastructure supported by a grant from the UK Medical Research Council (MR/L015080/1).

